# Inference of differential gene regulatory networks from gene expression data using boosted differential trees

**DOI:** 10.1101/2022.09.26.509450

**Authors:** Gihanna Galindez, Markus List, Jan Baumbach, David B. Blumenthal, Tim Kacprowski

**Affiliations:** Division Data Science in Biomedicine, Peter L. Reichertz Institute for Medical Informatics of Technische Universität Braunschweig and Hannover Medical School, Braunschweig, Germany; Braunschweig Integrated Centre of Systems Biology (BRICS), TU Braunschweig, Braunschweig, Germany; Chair of Experimental Bioinformatics, TUM School of Life Sciences, Technical University of Munich, Munich, Germany; Institute for Computational Systems Biology, University of Hamburg, Hamburg, Germany; Computational Biomedicine Lab, Department of Mathematics and Computer Science, University of Southern Denmark, Odense, Denmark; Department Artificial Intelligence in Biomedical Engineering (AIBE), Friedrich-Alexander-Universität Erlangen-Nürnberg (FAU), Erlangen, Germany

## Abstract

Diseases can be caused by molecular perturbations that induce specific changes in regulatory interactions and their coordinated expression, also referred to as network rewiring. However, the detection of complex changes in regulatory connections remains a challenging task and would benefit from the development of novel non-parametric approaches. We developed a new ensemble method called BoostDiff (boosted differential regression trees) to infer a differential network discriminating between two conditions. BoostDiff builds an adaptively boosted (AdaBoost) ensemble of differential trees with respect to a target condition. To build the differential trees, we propose differential variance improvement as a novel splitting criterion. Variable importance measures derived from the resulting models are used to reflect changes in gene expression predictability and to build the output differential networks. BoostDiff outperforms existing differential network methods on simulated data evaluated in two different complexity settings. We then demonstrate the power of our approach when applied to real transcriptomics data in COVID-19 and Crohn’s disease. BoostDiff identifies context-specific networks that are enriched with genes of known disease-relevant pathways and complements standard differential expression analyses. BoostDiff is available at https://github.com/gihannagalindez/boostdiff_inference.

**Author Summary:** Gene regulatory networks, which comprise the collection of regulatory relationships between transcription factors and their target genes, are important for controlling various molecular processes. Diseases can induce perturbations in normal gene co-expression patterns in these networks. Detecting differentially co-expressed or rewired edges between disease and healthy biological states can be thus useful for investigating the link between specific disease-associated molecular alterations and phenotype. We developed BoostDiff (boosted differential trees), an ensemble method to derive differential networks between two biological contexts. Our approach applies a boosting scheme using differential trees as base learner. A differential tree is a new tree structure that is built from two expression datasets using a splitting criterion called the differential variance improvement. The resulting BoostDiff model learns the most differentially predictive features which are then used to build the directed differential networks. BoostDiff outperforms other differential network methods on simulated data and outputs more biologically meaningful results when evaluated on real transcriptomics datasets. BoostDiff can be applied to gene expression data to reveal new disease mechanisms or identify potential therapeutic targets.

## 1. Introduction

Gene regulation is a fundamental biological process that underlies various cellular functions, including developmental, environmental, and disease contexts. The regulatory relationships in a biological sample can be represented by gene regulatory networks (GRNs), where two gene nodes with a regulatory relationship are connected by an edge [1]. GRN inference remains a challenging task because of the inherent complexity of transcriptional regulation, as well as the high dimensionality and noise in biological datasets. Furthermore, GRNs are dynamic and context-specific [2,3], i.e. some regulatory processes are active only in certain cell types, tissues, conditions, or in response to specific stimuli. Changes in these pairwise dependencies have been associated with the development of complex diseases [4]. Differential network analysis, which aims to detect altered connectivity between different conditions or disease states, has recently emerged as a powerful complement to standard differential expression (DE) analysis and is more suitable for detecting context-specific GRNs [4,5]. Exploring how GRN structures are rewired between two different states can reveal molecular mechanisms that drive disease development and progression and identify more relevant therapeutic targets.

Various approaches for deriving differential networks have been the focus of recent studies [6–8]. Representative methods are shown in Table 1. The z-score method performs Fisher transformation of Pearson’s correlation coefficients between two conditions. The resulting z-scores are modeled as a normal distribution, followed by a z-test to detect significant pairwise edges [9]. Diffcoex first builds an adjacency matrix and subsequently finds differentially co-expressed gene clusters using the topological overlap measure as a dissimilarity metric [10]. Another approach, the Gaussian graphical model (GGM)-based method, learns the differential network from conditional associations [11]. EBcoexpress relies on empirical Bayes’ estimation to estimate the posterior probability that an edge is differentially co-expressed [12,13].

**Table 1.** Overview of differential network methods used for comparison (adapted from Bhuva et al. [7]).

| Differential network method | Algorithmic approach | Test | Directionality | No. of conditions | Reference |
| --- | --- | --- | --- | --- | --- |
| BoostDiff | Tree-based | – | Yes | Two | This paper |
| z-score | Correlation-based | z-test | No | Two | [9] |
| EBcoexpress | Empirical Bayes + correlation | – | No | Two | [12] |
| Diffcoex | Correlation-based | Permutation test | No | Multiple | [10] |
| GGM-based | Gaussian graphical model + posterior odds | – | No | Two | [11] |
| chNet | Gaussian graphical model + differential expression analysis | t-test | No | Two | [14] |

The differential network methods described above measure linear relationships or rely on joint normality assumptions, which may not hold in practice [15]. In real biological datasets, complex, higher-order dependencies may be difficult to detect using correlation- or GGM-based methods. As discussed in a recent review, new methods for differential network analysis for non-Gaussian data are needed [15]. In this respect, tree-based strategies offer the advantage of more relaxed model assumptions. While examples such as GENIE3 and derived tools continue to be successfully applied in various biological settings [16,17], they cannot be used to compare different biological conditions.

We introduce BoostDiff, a non-parametric approach for reconstructing directed differential networks (Fig 1). We modified standard regression trees to identify gene pairs that show changes in regulatory dependencies between two biological conditions. To build the differential trees, we use a novel splitting criterion called the differential variance improvement (*DVI*), which measures the difference in predictive value of a feature on gene expression levels between two conditions. We demonstrate that boosting the differential trees with respect to samples belonging to a target condition is an important step for promoting condition specificity of the output networks. Tree-based variable importance measures can then be used to obtain a ranking of regulators.

**Fig. 1.**
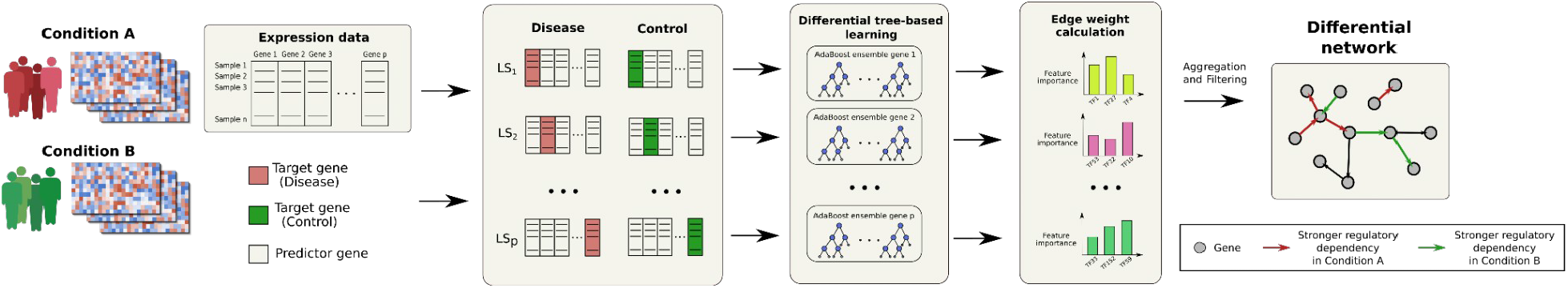
Overview of the BoostDiff algorithm. As input, we require two gene expression matrices corresponding to a target condition (e.g. disease) and a baseline condition (e.g. control). For each of *p* total genes, a learning subsample (LS) is drawn from the two datasets, after which an AdaBoost ensemble of differential trees is built to identify the features that are more predictive of the gene expression levels in the target condition. By setting a target condition, BoostDiff can be used to identify regulatory relationships that are more pronounced in condition A (e.g. disease state) and condition B (e.g. control/healthy), thereby providing a differential network capturing context-specific regulatory changes. In the overall workflow, the BoostDiff algorithm is run twice, one with condition A as target condition and subsequently with B as target condition. The results are then combined to obtain the final differential network. Most notably, while existing approaches aim for the reconstruction of whole genome-scale GRNs, BoostDiff concentrates on maximizing the precision for those parts of the regulatory network that actually predict the difference between the two phenotypes.

## 2. Methods

### 2.1 Overview of the differential network inference approach

The differential network inference problem can be decomposed into *p* independent regression subproblems, where *p* is the total number of genes in the expression data. Our strategy assumes that, in a given biological context, the expression level of a gene can be modeled as a function of the expression levels of other genes (Fig 1). This overall principle has been described in GENIE3 [16].

The crucial difference between BoostDiff and GENIE3 is that we simultaneously take into account two datasets for inferring a differential network. More precisely, our approach requires the availability of (1) gene expression data matrix 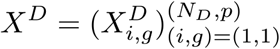 for *N*_*D*_ measurements from a disease condition and (2) the matrix 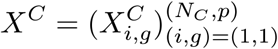 for *N*_*C*_ measurements from a control condition, both having *p* total genes (columns). The inference task can be viewed as a feature selection problem that aims to find features that are more predictive of expression levels in a target condition than in the baseline condition. In other words, differential network analysis is performed by solving the regression problem while taking into account information from two distinct labels. To achieve this, we employed the AdaBoost algorithm using differential trees as base learners to drive the improved prediction of expression levels in the target condition. The trained model provides a ranking of the edges by deriving a feature importance weight for each regulator.

A higher feature importance value means that the gene is more predictive and thus provides evidence of a stronger regulatory effect in one condition relative to the other. For each gene *g* = 1, … *p*, we define regression problems 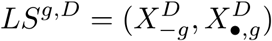 and 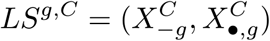. The design matrices 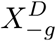 and 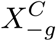 are obtained by deleting the *g*^*th*^ columns from *X*^*D*^ and *X*^*C*^, respectively, and the target variables are set to the deleted columns. The inference is performed as follows:

1. For *g* = 1, … *p*:
  a. Generate the learning samples of input-output pairs 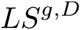 and 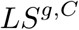 for gene *g*.
  b. Use a feature selection technique on 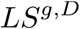 and 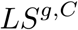 to calculate weights for all predictor genes except for itself. Here, an AdaBoost ensemble of differential trees is used as the feature selection technique.
2. Aggregate and sort the individual gene rankings to obtain a global ranking of all differential regulatory edges.

### 2.2 Growing a differential tree

In the following, we describe the steps to build a single differential tree, assuming we start with the learning samples 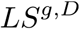 and 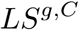 as input. A differential tree is built through binary recursive partitioning. The key difference to standard regression trees is that, to determine the features (i.e., genes) used for splitting the samples at the inner nodes of our trees, we use a novel split criterion called differential variance improvement (*DVI*) instead of variance reduction.

At each node of the differential tree, we maintain subsets *S*^*D*^ ⊆ 1, …, *N*_*D*_ and *S*^*C*^ ⊆ 1, …, *N*_*C*_ of the rows of 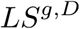 and 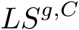 corresponding to the disease and control samples, respectively. Given a possible split feature (i.e., candidate predictor gene) *g′*, we define *DVI*(*g′*) as follows:

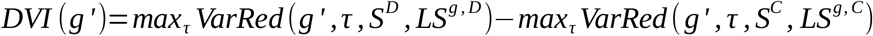

For fixed *g′* and splitting threshold *τ*, the variance reduction for the disease samples is given by:

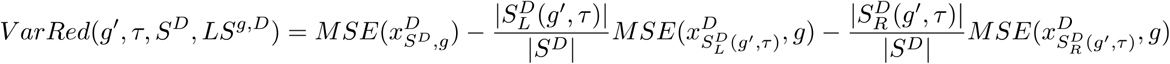

*MSE* is the mean squared error from the sample mean used as the impurity measure, 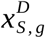 is the restriction of the target variable to the disease samples (rows) contained in a set of samples *S*, and 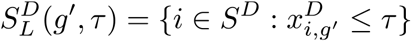 and 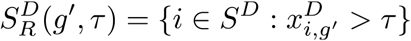 are the subsets of disease samples that fall to the left and right children of the candidate node, respectively.

Variance reduction for the control samples is defined analogously. A positive value of the *DVI* hence means that the gene *g′* is more predictive of *g*’s expression level in the disease condition than in the control condition, whereas a negative *DVI* value indicates that the opposite is the case.

Given training sets 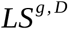 and 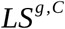 for the disease and control conditions, respectively, we construct a differential regression tree whose nodes are 5-tuples 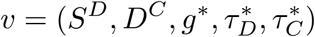, where *g*^*∗*^ is the split gene, 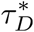 is the split threshold for the disease samples, and 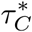 is the split threshold for the control samples. Note that we use two different thresholds, since using a single threshold for both conditions while optimizing the *DVI* will lead to a skewed expression distribution in each side of the split, with one side favoring disease samples and the other side favoring control samples. The construction is done as follows:

1. Initialize root as *r* = (*S*^*D*^ = 1, …, *N*_*D*_, *S*^*C*^ = 1, …, *N*_*C*_, •, •, •), where “ • ” is a placeholder for not yet defined split genes and thresholds.
2. Starting at the root, recursively construct a differential tree via binary partitioning as follows:
3. At the current node *v* = (*S*^*D*^, *S*^*C*^,•, •, •)of the tree under construction, do the following:
  a. If a suitable termination criterion (maximum depth or minimum number of target or baseline samples) has been reached or *max*_*g′*_ *DVI*(*g′*) ≤ 0, label *v* as leaf and traceback.
  b. Otherwise, set the node *v*’s split gene to *g*^*•*^ = *argmax*_*g′*_ *DVI*(*g′*), its disease threshold to 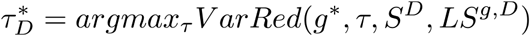, and its control threshold to 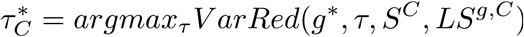.
  c. Initialize *v*’s left child as 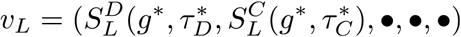 and its right child as 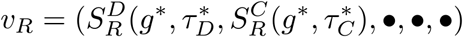 and continue with processing *v*_*L*_ and *v*_*R*_.

Ultimately, the differential tree learns a hypothesis *h*(*x*), → *y*, where *y* ∈ ℝ. In the regression trees described by Breiman [18], the prediction for a sample is determined by traversing the tree until a leaf node is reached. Here, we are more interested in predicting the expression values of the samples in the target condition; thus, prediction is performed only for target samples using the identified splitting thresholds 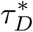. The final prediction is calculated as the expected value of the expression levels of the target samples assigned to the leaf nodes after fitting the differential tree.

### 2.3 Boosted differential trees

Inspired by GRNBoost2 [17], we implemented a boosting algorithm that derives a strong prediction model by sequentially training a pool of differential trees as weak learners. AdaBoost for regression is typically used for solving problems where the output is a continuous variable (i.e. expression levels) without explicitly considering the class of the samples. Here, we adapted the AdaBoost.R2 algorithm [19] to handle the regression problem given labels from two classes (i.e. conditions). Using the differential trees as base learners, the modified algorithm performs the boosting with respect to samples belonging to the specified target condition. The algorithm is described in detail in S1 Text. In this way, BoostDiff attempts to find a model that is more predictive of the target condition compared to the baseline condition. In each tree, only the target samples are re-weighted in subsequent boosting iterations, while samples from the baseline condition retain uniform weight. In particular, target samples that are more difficult to predict are selected with higher weights during the bootstrapping step and will always be compared to a uniform sample from the baseline condition. To avoid overfitting, we set a low number of trees and in practice find that 50 to 100 differential trees in the ensemble is sufficient for real datasets.

### 2.4 Variable importance measure

Tree-based methods allow for the calculation of a variable importance measure that can be used to rank the features according to their relevance for predicting the output. In GENIE3, the importance of a predictor gene *g′* is calculated as the sum of the variance reduction across all nodes where *g′* is used as the splitting feature, averaged over all trees in the ensemble. In the context of differential trees, we can derive a similar measure by considering the samples belonging to the target condition (i.e. disease samples). The importance attributed to a predictor gene *g′* can be calculated as the weighted variance reduction across *M* trees in the ensemble:

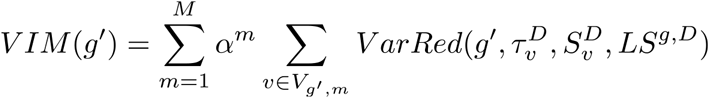

here *m* is the boosting iteration, *α*^*m*^ is the weight of the differential tree returned by AdaBoost, *V* _*g’, m*_ is the set of nodes in the tree where *g′* was used as the splitting feature, 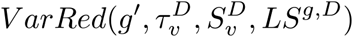 is the variance reduction given *g′*, the disease threshold 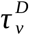, and the set of disease samples samples 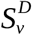 at node 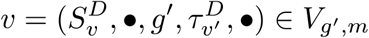 (see S1 Text).

Notably, because each node in a differential tree has two independent thresholds, interpreting the tree becomes more abstract with increasing depth. Boosting using shallow differential trees (e.g. differential tree stumps) thus favors greater interpretability of the variable importance measure.

### 2.4 Edge ranking and filtering from boosted differential trees

Each modified AdaBoost model yields a separate ranking of the regulators. However, simply ordering the regulatory links according to the weights leads to a bias for highly variable predictor genes. To avoid this, we first scale the expression levels of each target gene to unit variance, similarly implemented in GENIE3 [16].

Boosting with respect to a target condition does not necessarily produce a model that predicts a gene’s expression in the target condition better than its expression in the baseline condition. To illustrate, sample plots of the training progression are shown in S2 Fig. To restrict the results to differential edges, we recommend examining the distributions of the mean difference in prediction error. Sample distributions of these values from the simulated and real transcriptomics data are shown in S3 and S4 Figs, respectively. Based on these generated plots, users can filter for target genes with lower mean prediction error in the target condition than the baseline condition by applying a threshold. Alternatively, users can select the top edges with the lowest mean difference in prediction error or input a user-defined percentile. After filtering, the edges are re-ranked based on the variable importance measure used as the edge weight. The top *n* edges are then output as the final context-specific network.

## 3. Results and Discussion

### 3.1 Compared methods

To verify the condition specificity of the output networks, we first compared the boosted differential trees to a baseline random forest of differential trees, as well as two popular GRN inference methods, GENIE3 and ARACNE [20]. GENIE3 infers a network by building an ensemble of regression trees (i.e. random forest) [16]. It was run using the corresponding R package. ARACNE calculates the mutual information (MI) between all pairs of genes [20]. Afterwards, based on the data processing inequality (DPI) [21], it goes through all gene triplets and removes the edge with the weakest MI value. ARACNE was run using the implementation provided in the R package minet [22]. For both GENIE3 and ARACNE, only the disease expression matrix was used as input. For details, an AIMe report is available at https://aime.report/656I3Z/2 [23].

Next, we compared the performance of BoostDiff to other differential network methods. The benchmarking study conducted by Bhuva *et al*. indicated that the z-score method and EBcoexpress perform well in detecting differential edges compared to other methods [7]. Thus, we compared BoostDiff to z-score and EBcoexpress, as well as Diffcoex and a GGM-based method. Additionally, we run the more recently proposed chNet algorithm [14], which considers significant changes in both partial correlations of edges and differential expression. To facilitate comparability and given that only BoostDiff provides directionality information among the methods examined here, we converted directed edges to undirected edges [7].

### 3.2 Evaluation using simulated data

Gene expression data for disease and control conditions were simulated by adapting the SimulatorGRN approach [7], which simulates differential co-expression by knocking down nodes in the reference GRN by reducing their expression levels. In the original SimulatorGRN framework, a sample can have multiple genes knocked down, even though the evaluation considers each knockdown gene separately. To eliminate the confounding effect of additional knockdown genes in our experiments, we generated the expression data in the perturbed condition such that exactly one randomly selected input gene is knocked down. We evaluated the different tools based on two scenarios, namely, using networks with 150 nodes and 300 nodes, with 500 simulations per scenario. In each simulation, 100 samples were generated per condition. The final disease samples were those which have a gene knocked down, whereas the control samples are wild-type. We measured the performance of the algorithms with respect to the association network of the SimulatorGRN framework. The hyperparameter settings for generating the simulated data are shown in S1.

In all analyses on simulated data, all genes except for the target gene were considered as potential regulators. The z-score method, EBcoexpress, chNet, Diffcoex, and the GGM-based method were run with the default parameters. The parameters used for the random forest of differential trees and BoostDiff are provided in S2 Table. For the COVID-19 dataset, 50 trees were used, while 100 trees were used for the Crohn’s disease dataset because of the low sample size available for inference. For each simulation, we filtered for the target genes belonging to the 3rd percentile based on the mean difference in prediction errors (S3 and S4 Figs).

BoostDiff is designed to identify the predictive regulatory relationships that are more pronounced in a target condition relative to the baseline condition. Thus, to obtain a more complete differential network, the algorithm is run twice, once using the disease condition as the target condition (with control as the baseline condition) and another using the control condition as the target condition (with disease as baseline condition). In general, combining the two results performs better than the individual sub-analyses, indicating that each run can contribute meaningful edges to the output (S1 Fig). For subsequent comparisons with other inference methods, the combined results are presented.

The different tools have different statistical methods and cutoffs for determining the differentially coexpressed edges depending on how the algorithm works. To facilitate comparability, we show the top k predicted edges output by each method (except for chNet, wherein the number of predicted differential edges depends on the tuning parameter and is variable for each simulation; thus, extracting the top k edges cannot be consistently applied across simulations). For visualization, we show results based on the top 100 predicted genes output by each method. We report the performance using precision, recall, and F1 score as the evaluation metrics. Results were similar for varying cutoffs of k=50, 100, 150 and 200 (S6 Fig).

As expected, compared to GENIE3 and ARACNE, both of which infer a static network, BoostDiff can better identify the differential edges (Fig 2). The boosting scheme also performs significantly better than the random forest of differential trees. Importantly, BoostDiff outperforms the other differential network methods in all three metrics in both settings with 150 and 300 nodes.

**Fig. 2.**
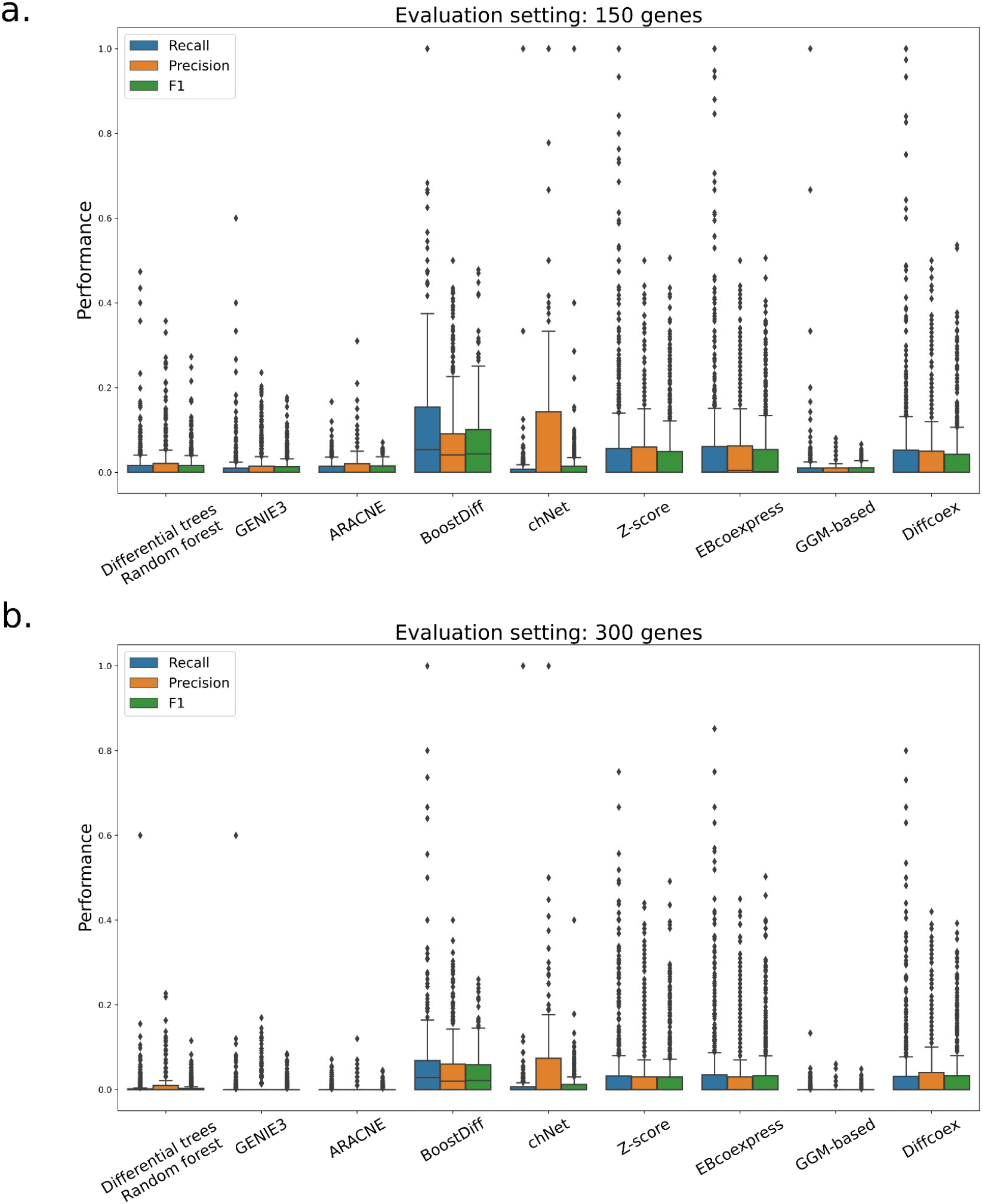
Performance of differential trees and boosted differential trees compared to the standard GRN inference methods and other differential network methods using simulated data comprising a) 150 genes and b) 300 genes. A total of 500 simulations were generated per evaluation setting. BoostDiff outperforms all other methods in both scenarios and can better identify the differentially co-expressed genes.

### 3.3 Evaluation using real datasets

We evaluated BoostDiff using a publicly available COVID-19 RNA-Seq dataset. Raw gene counts were downloaded from the Gene Expression Omnibus (GEO) database under the accession number GSE156063 [24]. We used data generated from nasal swab samples from COVID-19 (n=93) and uninfected patients (n=100). Count data were normalized using the DESeq2 package in R with the variance stabilizing transformation (vst) function. We also ran BoostDiff on a Crohn’s disease (CD) dataset. Normalized microarray data were downloaded from the GEO database under the accession GSE126124 [25] using data generated from colon biopsies of individuals with Crohn’s disease (n=37) and healthy controls (n=19). Illumina IDs were converted to HGNC symbols using the R package biomaRt [26]. Expression levels corresponding to probes mapped to the same gene symbol were averaged. Differentially expressed genes (DEGs) were obtained using DESeq2 for the COVID-19 dataset and using limma for the Crohn’s disease dataset [27,28].

The z-score method and EBcoexpress were run with default parameters. The parameters used for the BoostDiffs run are provided in S2 Table. For the Crohn’s disease dataset, a higher number of trees (100 estimators) were used because of the lower number of samples available for inference. The list of human transcription factors downloaded from http://humantfs.ccbr.utoronto.ca/ were used as the candidate regulators [29]. For the COVID-19 dataset, data were already normalized with the vst function, so we set normalize=False. All the outputs from the different methods were filtered for the top 1000 edges (except for chNet). For BoostDiff, the final network thus comprised the top 500 edges from the run where the disease condition was set as the target condition, and the top 500 edges from the run where the control condition was set as the target condition. Genes whose mean difference in prediction error of the models were more extreme than the threshold identified from the 3rd percentiles of the distributions were retained. The enrichr module of the gseapy package was used to identify enriched KEGG pathways in the output networks [30,31]. The Louvain community detection algorithm was applied using the python-louvain package (https://github.com/taynaud/python-louvain).

#### 3.3.1 COVID-19

The differential network output by BoostDiff is enriched with pathways that are consistent with known COVID-19 pathophysiology. In addition to various pathogenic infections, such as “Herpes simplex I infection”, “Human T-cell leukemia virus,” “Influenza A,” “Epstein-Barr virus infection”, and “Measles”, the output network was significantly enriched in COVID-19-relevant pathways, such as “Coronavirus disease,” “Th17 cell differentiation,” “IL-17 signaling pathway,” “NF-kappa B signaling pathway,” “NOD-like receptor pathway,” “Toll-like receptor signaling pathway,” and “TNF signaling pathway” (Fig 3). Toll-like receptors (TLRs) are involved in the innate immunity and function in pathogen recognition and cytokine regulation. Infection by SARS-CoV-2 particularly triggers TLR2. TNF is a key cytokine that drives inflammatory macrophage phenotype and tissue damage in severe COVID-19 [32]. The NF-κB pathway activation contributes to the cytokine storm that affects critically ill patients. Both NF-κB and TNF signaling have been proposed as therapeutic targets to prevent organ damage in COVID-19 [33]. Viral infections activate NOD-like receptors, which lead to inflammasome assembly [34]. Th17 signaling participates in the cytokine response characteristic of the “cytokine storm” and leads to the production of IL-17 [35,36]. Th17 cells were found to undergo more clonal expansion in the lungs of severe COVID-19 patients [37]. Imbalance in the Th1 and Th2 signaling has also been associated with COVID-19 mortality risk [38]. Examining the differential edges when considering the two sub-analyses separately shows generally similar results, indicating enrichment of infection related pathways (S10 Fig). The differential network output by the z-score method did not show the enrichment of COVID-19-specific pathways (S8 Fig), whereas all edges in the EBcoexpress output showed zero posterior probabilities.

**Fig. 3.**
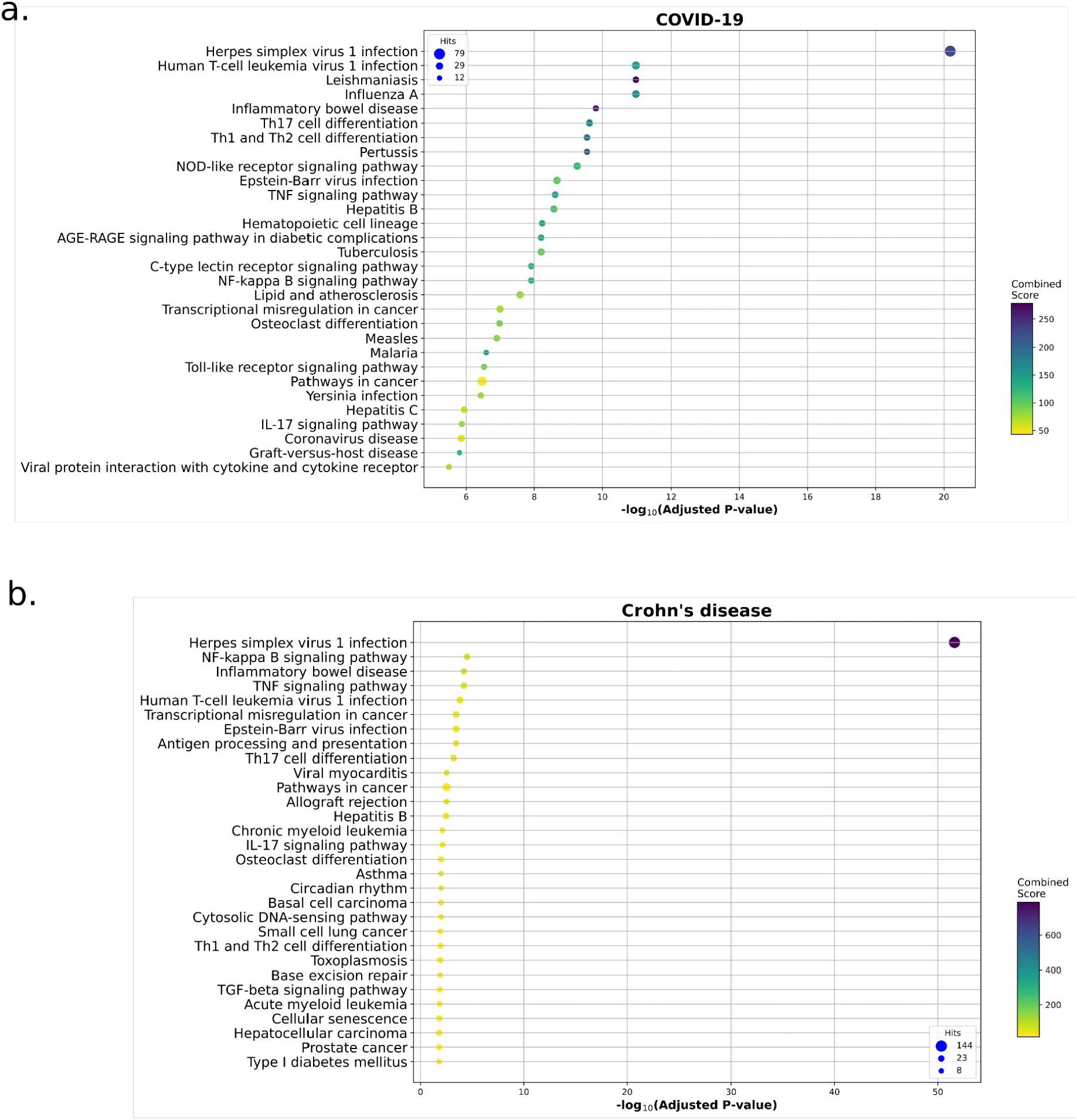
Enriched KEGG pathways in the network inferred by BoostDiff for the a) COVID-19 dataset and b) Crohn’s disease dataset.

We also compared the BoostDiff network to the list of DEGs. While the overlap between the differential network and DEGs is significant, it is quite low (Jaccard similarity=0.125). Further, removing the DEGs from the genes in the differential network retained the enrichment of COVID-19-related pathways (S11 Fig), indicating that these dysregulated genes identified by BoostDiff are missed by standard DE analysis. Performing enrichment analysis separately for the targets and regulators of the predicted edges showed similar results, demonstrating the effectiveness of the feature selection approach (S12 Fig).

To further examine the differential network output by BoostDiff, we applied the Louvain community detection algorithm [39], which produced a total of 84 modules. We identified a dysregulated cluster comprising 59 genes that showed enrichment in the terms “Chemokine signaling pathway,” “Viral protein interaction with cytokine and cytokine receptor”, “Coronavirus disease,” “Toll-like receptor pathway,” and “Th1 and Th2 cell differentiation” (Fig 4). Notable coronavirus disease-related genes in this module include *CXCL10, DDX58, STAT1, STAT2, EIF2AK2*, and *ISG15*. Other additionally known genes involved in pathogen response include *IFIT1, IFIT2, IFIT3, CXCL11, CXCL9* and *CCR1*. Chemokines are produced in response to a range of viral infections. In COVID-19, chemokine signaling has been linked to acute respiratory distress syndrome [40]. *DDX58* (RIG-1) is involved in the production of interferons in response to COVID-19 [41]. Interferon signaling mediated by *STAT1* and *STAT2* is a key antiviral defense mechanism. The chemokines *CXCL9, CXCL10* and *CXCL11* are known to be upregulated in the COVID-19 response [42]. *EIF2AK2* is an interferon-induced protein kinase that plays a role in inhibiting viral replication [43]. *IFIT1, IFIT2*, and *IFIT3* form a functional complex and participate in interferon-induced broad viral response[44,45]. *ISG15* is a ubiquitin-like protein whose activation triggers the release of various pro-inflammatory cytokines and chemokines [46]. Polymorphisms in *HLA-DRB1* have been reported in severe COVID-19 patients [47]. The expression of the antigen presentation gene *HLA-DQA2* has been reported to be downregulated in severe cases [48]. Based on these results, further experimental validation in this module would be of interest to uncover a more detailed mechanistic understanding of COVID-19 disease pathogenesis.

**Fig. 4.**
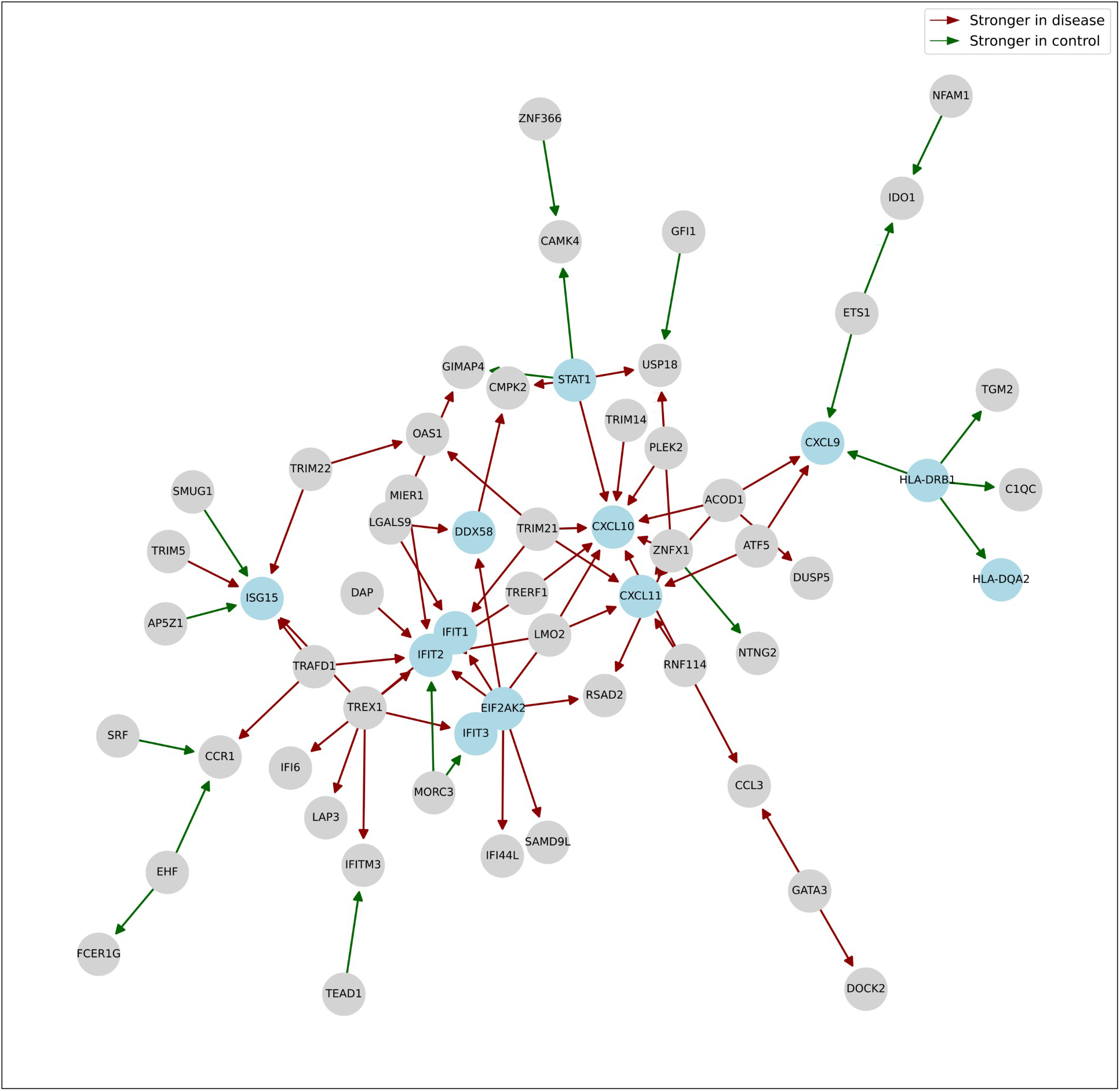
Dysregulated module identified from the COVID-19 differential network inferred by BoostDiff using the Louvain algorithm. Notable genes in the module include *CLCL9, CXCL10, CXCL11, DDX58, STAT1, IFIT1, IFIT2, IFIT3, EIF2AK2, HLA-DRB1*, and *HLA-DQA2*, which are highlighted in blue.

#### 3.3.2 Crohn’s disease

Crohn’s disease (CD) and ulcerative colitis (UC) are the two main types of inflammatory bowel diseases (IBDs). CD is an autoimmune disease characterized by chronic inflammation of the gastrointestinal tract and impaired intestinal barrier function. IBDs are thought to be caused by a complex interplay between the gut microbiome, the host immune system, and the environment. Using a Crohn’s disease dataset derived from CD patients and healthy controls, we derived differential networks using the z-score-based method and EBCoexpress. Although sample sizes were relatively low for this dataset, the CD-specific differential network output by BoostDiff was enriched in CD-relevant pathways, including “Inflammatory bowel disease,” “Th17 cell differentiation,” “IL-17 signaling pathway”, “NF-κB signaling,” “Antigen processing and presentation”, “TGF-β pathway,” and “TNF signaling pathway” (Fig 4 and S2 Table). Toll-like receptors (TLRs) play a role in host defense and homeostasis by acting as sensors of microbial pathogens. IBD has been associated with abnormal gut microbiota composition and TLR overstimulation, which in turn promotes NF-κB signaling and downstream inflammatory responses [49]. TGF-β signaling plays an immunosuppressive role in mucosal inflammation, and impaired signaling can lead to intestinal fibrosis [50,51]. NF-κB is a transcription factor that functions in maintaining intestinal homeostasis, and dysregulation of the NF-κB pathway leads to sustained inflammatory state characteristic of IBD patients [52]. NF-κB signaling activation has been associated with more severe clinical manifestations in CD patients [52,53]. The Th17 subset of CD4+ T cells have well recognized roles in IBD pathogenesis. In CD, IL-17 signaling mediates the activation of Th17 cells, which further drive pro-inflammatory cascades via the production of IL-21, IL-22, IFN-γ and TNF [54]. The differential edges obtained from the sub-analysis where the control state was used as the target condition showed enrichment of further CD-relevant pathways (S13 Fig). Such cases may reveal more subtle differences in terms of differential predictivity of expression in two different disease states and motivate more refined downstream analysis by independently examining the results from the two sub-analyses.

Notably, the z-score method, while based on the correlation measure, did not return strong enrichment of disease-relevant pathways compared to BoostDiff. The z-score network was enriched in only one term, “Tryptophan metabolism.” While the output of EBcoexpress also identified the enrichment of several inflammatory pathways (S9 Fig), the differential network output by BoostDiff showed stronger enrichment based on the p-values. EBcoexpress on the full dataset took more than two weeks, whereas BoostDiff took less than one day (S2 and S3 Tables), thus limiting the applicability of EBcoexpress on real transcriptomics datasets. The parallelization of BoostDiff allows for more reasonable runtimes.

Differential expression analysis of the CD data identified ten DEGs, out of which only one was also present in the differential network identified by BoostDiff; consequently, enrichment results after removal of DEGs were similar to the original network (S14 Fig). We further performed enrichment analysis separately on the targets and regulators of the directed edges output by BoostDiff. As shown in S15 Fig, both the list of regulators and the list of targets from the differential edges were enriched in pathways related to Crohn’s disease, demonstrating the value of the *DVI*-based feature selection approach.

We applied the Louvain algorithm on the differential network output by BoostDiff, which identified a total of 326 modules. One interesting dysregulated module was enriched in multiple autoimmunity-related terms, including “allograft rejection,” “graft-versus-host disease,” and “autoimmune thyroid disease” (Fig 5). Notable genes in the module include *HLA-A, HLA-B, HLA-G, HLA-H*, and *HLA-J*. The human leukocyte antigen (HLA) is a genomic region that has been genetically linked to the susceptibility to autoimmune diseases and IBD [55]. The involvement of *HLA-G* in various autoimmune diseases, including UC and CD, are well documented [56]. The associations of *HLA-A, HLA-B, HLA-G, HLA-H*, and *HLA-J* with CD have been previously reported in eQTL and genome-wide association studies [57]. *TRIM21* (Ro52) has also been implicated in various autoimmune conditions [58]. In IBD, *TRIM21* is involved in regulating Th1/Th17 cell differentiation and mucosal inflammation [59]. *E2F2* belongs to the E2 family of transcription factors that plays a role in cell differentiation. *E2F2* expression in the colon is dysregulated in CD patients [60].

**Fig. 5.**
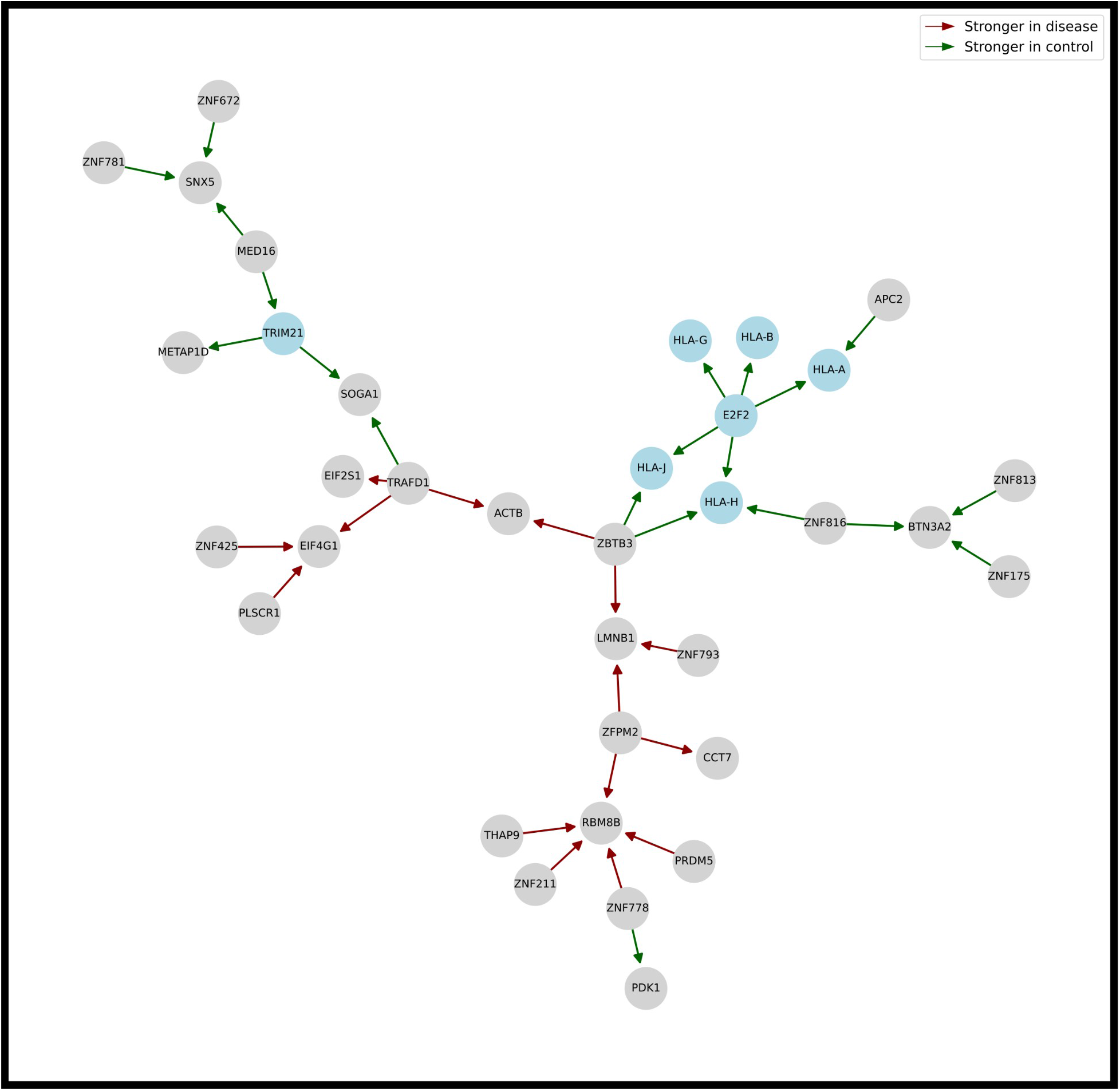
Dysregulated Louvain module identified from the Crohn’s disease differential network output by BoostDiff. Notable genes, namely, *HLA-A, HLA-B, HLA-G, HLA-H, HLA-J, TRIM21*, and *E2F2*, are highlighted in blue.

#### 3.3.3 Correlation distributions

We also examined the Pearson correlations of the top edges from the differential networks identified by BoostDiff using the original expression data. This procedure was performed separately for the results of the two sub-analyses, namely, when the disease condition is used as the target condition, and when the control condition is used as the target condition. As shown in Fig 6, for the same edges, we observe a unimodal distribution of correlation values in the non-target condition and a bimodal distribution where BoostDiff identified stronger associations in the target condition, where strong positive correlation values suggest activating regulator-target relationship in the target condition, while negative values indicate inhibitory relationships. These results are consistent with the goal of identifying differential co-expression between genes. This striking observation cannot be reproduced when compared to all pairwise edges from the list of DEGs or randomly selected edges. Differential edges in either condition tend to have highly correlated expression levels, indicating dysregulation based on disease status.

**Fig. 6.**
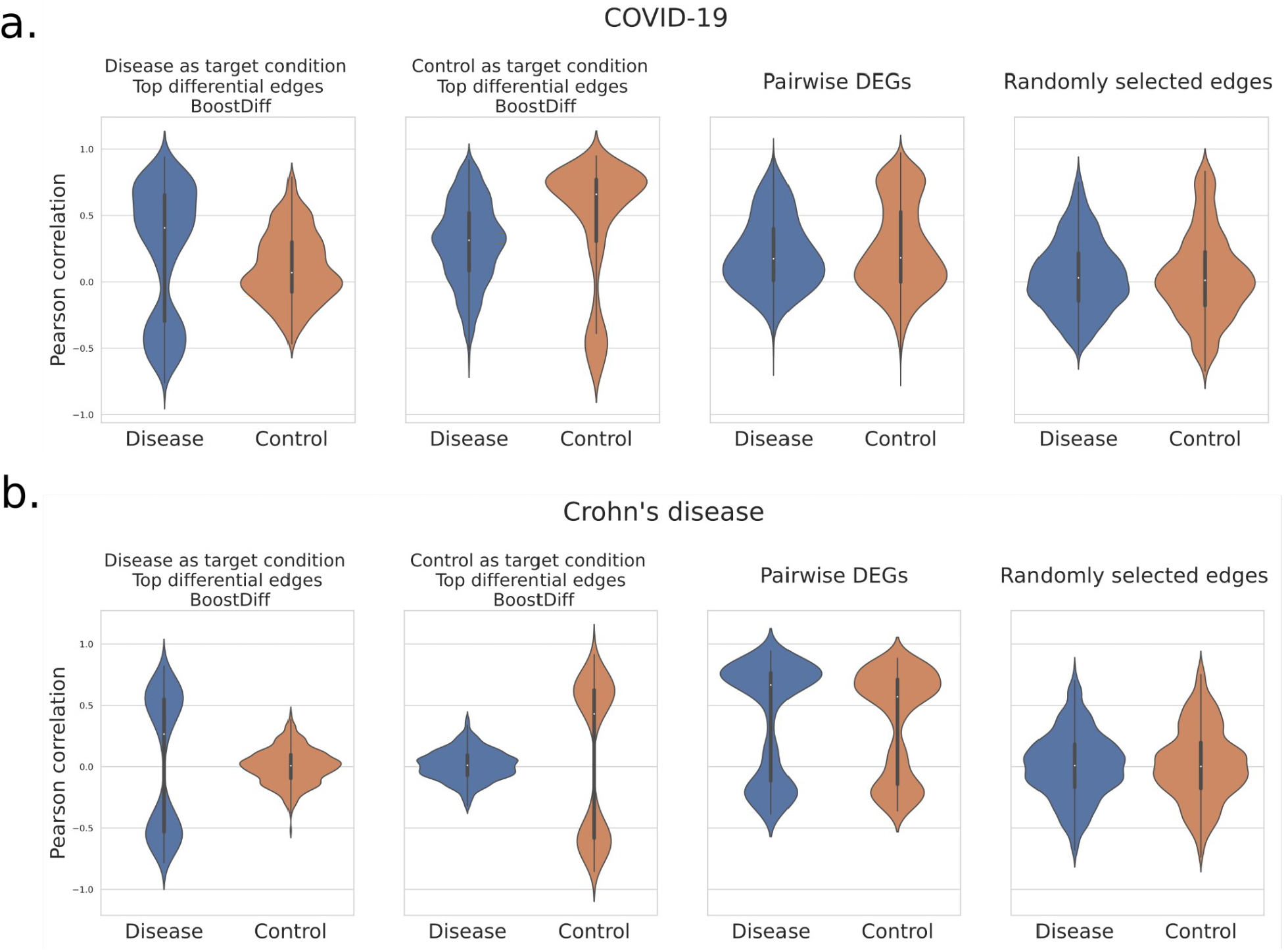
Violin plots showing that the top 500 edges in the differential network predicted by BoostDiff tend to exhibit changes in correlation distributions between the disease and control expression data, indicating dysregulation in pairwise relationships. Correlations between predicted differential edges are compared to correlations between all pairwise combinations of DEGs, as well as randomly selected edges. Results are shown for a) the COVID-19 RNA-Seq dataset b) the Crohn’s disease microarray dataset.

## 4. Conclusions

Gene regulation is a complex process that changes under different biological contexts. Differential network biology explores the rewiring of these regulatory interaction landscapes that are fundamentally distinct from the static networks that are inferred in most standard GRN inference methods [3]. By additionally considering the regulatory dependencies from a baseline condition, we can uncover a more refined picture underlying the molecular processes that are perturbed in a condition of interest, such as disease.

Inference of networks from biological expression data is a challenging task. The novelty of BoostDiff is twofold: 1) We employ differential variance improvement as the splitting measure in a tree-based algorithm that can explicitly compare two datasets with a continuous output variable; 2) BoostDiff adapts the AdaBoost algorithm to use differential trees as the base learner. Boosting the differential trees with respect to samples belonging to the target condition is a crucial step that significantly improves the detection of differential edges.

BoostDiff outperformed existing differential network methods on simulated data and can better handle the simulated datasets with higher dimensionality. BoostDiff yields biologically meaningful results and is more practically applicable on real-world transcriptomics datasets. We showed that the differential networks inferred by BoostDiff are consistent with the known pathophysiology of COVID-19 and Crohn’s disease. The performance of BoostDiff can be attributed to the tree-based nature of the algorithm, which performs inference of differential networks without assuming parametric distributions of gene expression. In particular, BoostDiff has more relaxed model assumptions and can better capture complex changes in gene dependencies in biological contexts, which could be missed by tools that employ correlation-based measures. BoostDiff is also scalable since it builds one model for each gene and can hence easily be parallelized.

Nevertheless, our method has several limitations. First, BoostDiff can only compare two conditions at a time. Moreover, BoostDiff is similar to GENIE3 in that it does not perform statistical testing. Instead, scores are assigned to individual edges by calculating tree-based variable importance measures; thus, only the ranking of the edge weights is considered. Further, the AdaBoost algorithm can be prone to overfitting, although this can be avoided by setting a low number of base differential trees.

The application of BoostDiff is not limited to gene expression data; the proposed feature selection approach can be generalized to other omics datasets. For instance, BoostDiff can be applied to proteomics or metabolomics data that aim to detect changes in dependencies of proteins or metabolites. Moreover, the simple but effective strategy implemented in BoostDiff is an algorithmic advancement that can be further extended to other problems that aim to extract differentially predictive features. Adapting BoostDiff for analyzing time-series datasets is also a promising research direction.

## Acknowledgements

We are grateful to Julian Beier for technical support.

## Author contributions

Writing – Original Draft Preparation: GG. Methodology: GG, DBB, TK. Software: GG. Supervision: DBB, TK. Conceptualization: ML, JB, DBB, TK.Writing – Review & Editing: ML, JB, DBB, TK.

## Data Availability

Transcriptomics data used for biological evaluation can be downloaded from Gene Expression Omnibus under the accession numbers GSE156063 and GSE126124. Our BoostDiff implementation is available on GitHub: https://github.com/gihannagalindez/boostdiff_inference.

